# Extraordinary adaptations: Functional and evolutionary synergy of trait components can explain the existence of leaf masquerade

**DOI:** 10.1101/2025.04.05.647375

**Authors:** J. Benito Wainwright, Charlotte E. J. Rolfe, Graeme D. Ruxton, Nathan W. Bailey

## Abstract

One of the most enduring mysteries in biology concerns the evolution of complex adaptations made up of interacting component traits. When these component traits lack an obvious adaptive function in isolation from one another, their origin requires either non-adaptive, intermediate evolutionary steps or their simultaneous, synergistic evolution. We tested these alternatives using the powerful but accessible example of leaf masquerade in katydids, where in some species, highly modified wings strikingly mimic vegetation to avoid predator recognition. Combining a field predation experiment with a phylogenetic comparative analysis of wing morphology in 51 Neotropical katydid species, we show that colour and shape synergistically interact to enhance survival in the wild, and modifications in both traits evolved concurrently during diversification of this clade. Our findings identify the functionality of masquerade camouflage in the wild and highlight how synergy between individual traits fosters evolution of extraordinarily specialised adaptations.

## Introduction

Understanding the evolutionary forces that shape phenotypic complexity is a fundamental component to all aspects of organismal biology. The degree of complexity varies widely across different traits. Composite adaptations, such as vertebrate eyes and bird wings, rely on the interaction of independent component traits to acquire an emergent synergistic function [1-3]. The evolutionary origin of such extreme adaptive complexity has been a persisting question since the birth of evolutionary biology, particularly when component traits do not appear to enhance fitness in isolation from one another [4,5]. For the springboard prey-trapping mechanism of *Nepenthes* carnivorous pitcher plants, it has recently been demonstrated that component traits arose independently, implying that complex composite adaptations can evolve through non-adaptive, intermediate evolutionary stages [3]. However, this route is not inevitable for other composite traits. An alternative scenario is that composite traits evolve when directional selection acts on individual components simultaneously [4]. In this case, individual components of composite traits should show a high degree of evolutionary correlation inconsistent with sequential acquisition through non-adaptive, intermediate stages. Historically, despite the ubiquity of complex traits, a lack of suitable examples for both comparative study and experimental manipulations has made it difficult to empirically measure their emergent fitness benefits under ecologically relevant conditions and dissect evolutionary pathways leading to their evolution in natural systems.

Leaf masquerade provides an accessible and promising example in which to evaluate these alternatives. Masquerade camouflage is the resemblance of organisms to objects in the environment such as leaves, twigs, stones, and bird droppings [6-9], causing predator misclassification independently of the background upon which they’re viewed [10]. Thus, masquerade provides protection without concealment. In some katydid lineages (bush crickets; Orthoptera: Tettigoniidae; Figure 1A), the tegmina (sclerotized forewings) have repeatedly evolved an extraordinary likeness to the shape and colouration of plant leaves from a non-leaf-like ancestor [11-13]. However, it is not known how the evolution of these features was coordinated to generate masquerade. It seems intuitive to suppose that a close resemblance in shape or colouration would provide little protective value from masquerade without the other. Nevertheless, katydid species appear to exhibit continuous variation in their degree of ‘leafiness’, which provides an opportunity to evaluate the functional and evolutionary explanations of this trait using experimental and comparative techniques.

**Figure 1.**
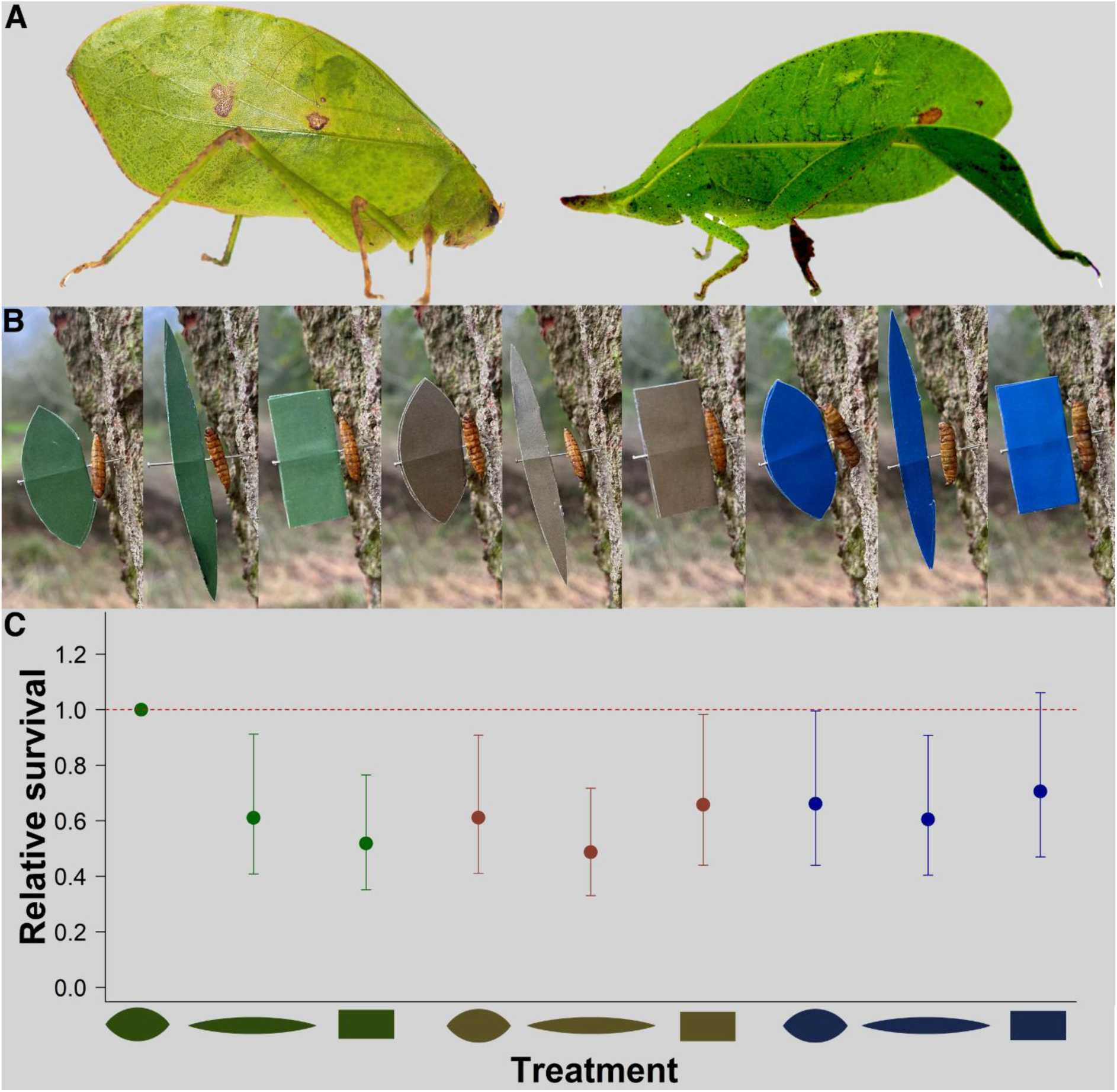
Functional synergy underlies effective leaf masquerade in the wild. (A) Photographs of *Itarissa costaricensis* (left) and *Aegmia maculofolia* (right), leaf-masquerading katydid species (Orthoptera: Tettigoniidae) found at the field site in Panama. © Photos courtesy of Dr Hannah ter Hofstede and Dr Laurel Symes. (B) All experimental colour x shape treatment combinations *in situ*, pinned to tree bark with a mealworm (*Tenebrio molitor* larvae) as bait. From left to right: ‘green oval’, ‘green elongated oval’, ‘green rectangle’, ‘brown oval’, ‘brown elongated oval’, ‘brown rectangle’, ‘blue oval’, ‘blue elongated rectangle’, ‘blue rectangle’. (C) Odds of targets being predated, derived from a fitted Cox survival model on all colour x shape treatment combinations (*N* = 1,296), relative to the ‘green oval’ treatment, which had the highest survival rate; this model is for illustrative purposes only. Red dashed line serves as a reference for comparison and error bars represent 95% confidence intervals.

Using artificial leaf-mimicking prey and naïve, free-living avian predators, we first tested how interactions among the component traits comprising leaf masquerade affect predator perception in the wild. Next, to assess how real leaf-mimicking species have evolved, we conducted phylogenetic comparative analysis on the tegmen morphology of 51 Neotropical katydid species and integrated this with measures of human perceptions of ‘leafiness’. Our data allow us to disentangle the evolutionary origins of leaf masquerade and shows how functional and evolutionary synergy can explain the existence of complex morphological ‘design’ in the natural world.

## Results and discussion

### Colour and shape non-additively enhance leaf masquerade in the wild

To determine how leaf masquerade improves individual fitness under ecologically relevant conditions, we exposed wild avian predators to artificial wing targets varying in colour to mimic naturally occurring green leaves, the brown bark substrate upon which they were pinned, or an unnatural blue control. Doing so meant we could confirm that any survival benefits incurred by these stimuli were a result of misclassification of the targets and not due to either enhanced crypsis of the targets against their background or to neophobic responses. Models of each colour also varied in shape to mimic an oval leaf, an elongate leaf, or a rectangular control (Figure 1B), and were designed based on morphological measurements taken from the tegmina of an existing leaf-mimicking katydid species (See Materials and methods). To minimise potential effects of learning, memory and other cognitive features that may have arisen to improve foraging efficacy in bird communities where leaf-masquerading species are native [14,15], we performed the experiment in a locality where avian predators do not routinely encounter leaf-masquerading insects. We found that the specific combination of green colour and oval shape increased fitness, rather than each trait improving survival in isolation (Figure 1C; Figure S1).

The interaction between colour and shape predicted survival rate (mixed-model Cox regression: χ^2^_4_ = 9.882, p = 0.043), but neither factor was significant as a main effect (mixed-model Cox regression: colour, χ^2^_2_ = 2.147, p = 0.342; shape, χ^2^_2_ = 5.878, p = 0.053). Furthermore, colour affected survival rate only for oval-shaped targets (mixed-model Cox regression: oval, χ^2^_2_ = 7.143, p = 0.028; elongated oval, χ^2^_2_ = 2.058, p = 0.357; rectangle, χ^2^_2_ = 2.814, p = 0.245; Figure 1C; see Table S1 for post-hoc tests). The ‘green oval’ targets did not match the background colour, and their shape were specifically designed to closely resemble that of an existing, but not local, leaf-masquerading katydid species, and likely other locally abundant leaves at the study site. Therefore, their increased survival compared to targets with brown, background-matching colouration and those with atypical leaf shapes strongly suggests that their survival advantage resulted from predators misclassifying them. This field experiment thus confirms an assumption central to the definition of masquerade that distinguishes it from other forms of camouflage [8,9,16].

### Composite trait evolution in Neotropical leaf-masquerading katydids

Our field predation experiment suggests that leaf masquerade is a composite trait reliant on the synergistic effects of colour and shape for its adaptive function of causing predator misclassification. To assess whether colour and shape evolved synergistically in real leaf-masquerading species, we studied the tegmina of 270 wild-caught individuals of 51 sympatric katydid species from a diverse Panamanian rainforest community (Figure 2A). The colouration of each species was classified (and independently verified; see Materials and methods) as being green or brown in colour and pigmented or not (Figure 2B; katydid tegmina can be translucent or opaquely pigmented irrespective of their general colour, with the former being the most likely ancestral state for Tettigoniidae). Our particular interest was in wings that were green *and* pigmented, since green targets provided better masquerade in the avian predation experiment above. All studied katydid species were cryptic in colouration and available data suggests they occupy similar rainforest microhabitats [17]. Therefore, shifts in colour and pigmentation do not offer any obvious additional protective value other than for the purpose of leaf masquerade. Shape was quantified by measuring the tegmen aspect ratio (maximum length / maximum width; see Materials and Methods). Thus, species with wider, more rounded, “leaf-like” wings had lower aspect ratios [18] (Figure 2C). The ecological relevance of our shape treatments in the avian predation experiment was supported by the observation that the species with the lowest aspect ratio (*Aegimia maculofolia,* aspect ratio = 2.177) closely matched that of the ‘oval’ treatment in our field experiment, while the species with the highest aspect ratio (*Caulopsis micropora,* aspect ratio = 8.841) closely matched that of the ‘elongated oval’ treatment. Expanding upon previous categorisations of Neotropical katydids [12], two ‘human leafiness metrics’ were obtained by asking 53 naïve human participants to both classify (presence vs. absence) and score (out of 10) leafiness of the 51 species’ tegmina via an online survey (see Materials and methods). Using a recent molecular phylogeny [19], we then reconstructed the evolution of leafiness and its shape and colour component traits.

**Figure 2.**
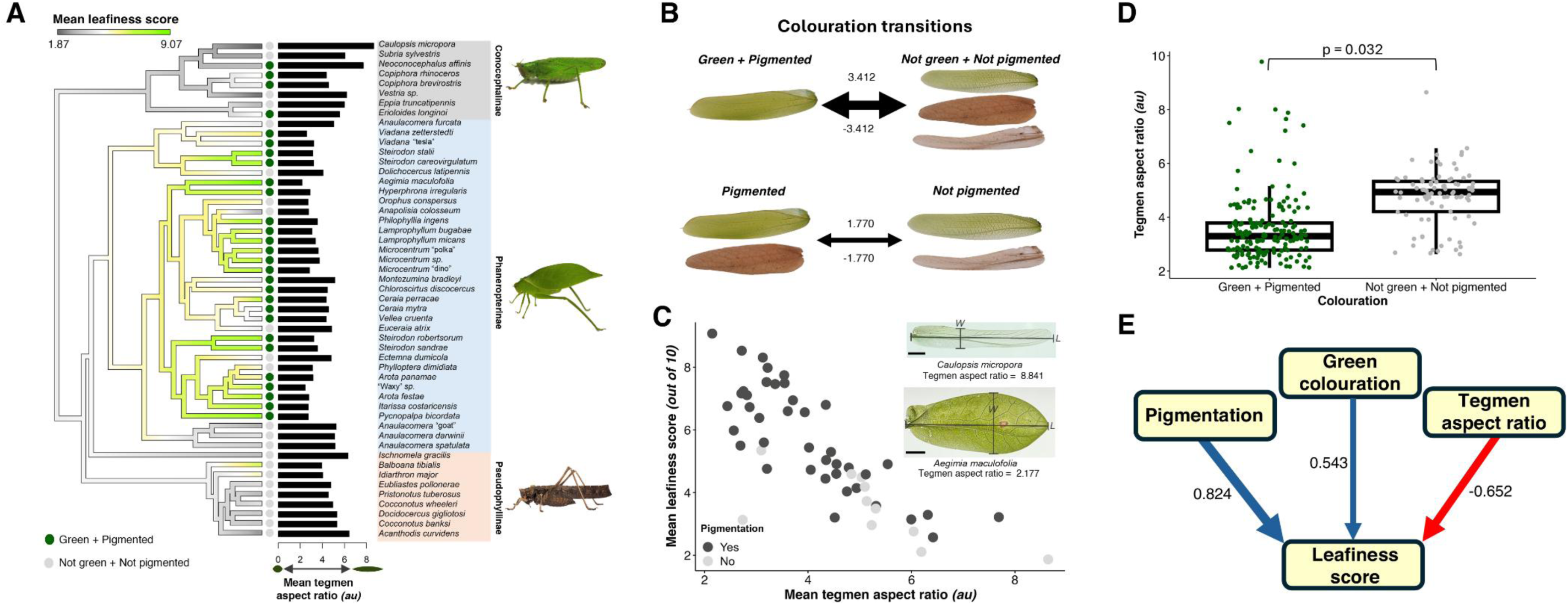
Synergistic evolution explains the origins of leaf masquerade in katydids. (A) A continuous trait plot reconstructing patterns of human perceived ‘leafiness’ (‘mean leafiness score’) per species across a pruned molecular phylogeny of 51 katydid species [19]. The presence (dark green) and absence (grey) of green pigmentation is shown at the tips, and the mean tegmen aspect ratio (au) per species is indicated by black horizontal bars. Exemplars from each subfamily (highlighted blocks) are shown on the right. © Photos courtesy of Ciara Kernan, Dr Laurel Symes and Dr Hannah ter Hofstede. (B) Transition rates between colouration states (above: green pigmentation, below: pigmentation). Arrow thickness is scaled by the transition rate value which is also given beside each arrow. (C) Non-additive effects of tegmen pigmentation (presence: dark grey, absence: light grey) and mean tegmen aspect ratio on the mean leafiness score per species (*N =* 51). The inset shows mounted tegmina of the species with the lowest (*Caulopsis micropora*) and highest (*Aegimia maculofolia*) mean aspect ratio, with the measured minor (*W*) and major (*L*) axis labelled for each. Scale bars = 5 mm. (D) Evolutionary association between tegmen aspect ratio (*N* = 270) and the presence and absence of green tegmen pigmentation across the 51 katydid species. Thick bars indicate medians, boxes indicate interquartile ranges, and whiskers show values within 1.5 interquartile ranges. (E) Phylogenetic pathway analysis illustrating the directional and simultaneous evolutionary acquisition of pigmentation, green colouration, and reduced aspect ratio during the evolution of leaf masquerade. Human perceived ‘leafiness score’ (out of 10) per species is the response variable. Arrows indicate the direction of the interaction, and values adjacent to these arrows are respective pathway coefficients indicating the strength of the estimated relationship. Positive coefficients (blue arrows) indicate a transition from absence to presence in the parent trait (tegmen pigmentation and green colouration) whereas negative coefficients (red arrows) indicate a decrease in the parent trait (mean tegmen aspect ratio).

We found that the evolution of leaf masquerade is under strong phylogenetic constraint. Both green pigmentation and shape show a strong phylogenetic signal (green pigmentation, χ^2^ = 0.000, λ = 1.000, p = 1.000; mean aspect ratio, χ^2^ = 13.445, λ = 1.061, p < 0.001) and transition rates between presence and absence of green pigmentation are equivalent (Figure 2B; Figure S2; Table S2,S3). The most likely pattern of aspect ratio evolution followed a Brownian motion (BM) model (BM vs Ornstein-Uhlenbeck: ΔAIC = -2.261, BM vs early burst ΔAIC = -1.710; Figure S2). Modelling the human leafiness metrics in the same way showed a similarly high phylogenetic signal, with a BM model being best fitting (Table S3). These results indicate a strong degree of evolutionary inertia, potentially limiting the axes of adaptive morphological evolution among leaf-masquerading species [20]. Consistent with previous work, ancestral state reconstructions support multiple independent elaborations and reductions of ‘leafiness’ and both its component traits across the 51 studied species [11] (Figure 2A).

### Colouration and shape have non-additive effects on human-perceived ‘leafiness’

In the face of this phylogenetic constraint, we sought to test whether the functional synergy of colouration and shape, indicated by our predation experiment, is concordant with human perceptions of leafiness. While humans are not natural predators of Neotropical katydids, many insectivorous New World primates are [21-23], and behavioural experiments have suggested conserved mechanisms underlying object recognition across birds and primates [24-26]. Because tegmina were presented to human participants against a white matte background (see Materials and methods), there is no biological reason why green tegmina were more likely to be classified as leaf-like over brown tegmina. Therefore, accounting for phylogenetic effects, we tested whether human perceptions of leaf masquerade depended on the interaction between shape and the presence or absence of notable tegmen pigmentation, independent of colour.

When individual participant data were analysed using phylogenetic generalized linear mixed models, the interaction between mean aspect ratio and pigmentation predicted the human leafiness score (MCMCglmm: P-mean = -4.696, 95% CI = -9.476 - -0.309, P_MCMC_ = 0.047; Figure 2A,C; Table S4). When species were subset by pigmentation, participants only awarded high leafiness scores to low aspect ratio tegmina if they were also pigmented, echoing results from the avian predation experiment (MCMCglmm: pigmented, P-mean = -8.576, 95% CI = -12.175 – -5.317, P_MCMC_ < 0.001; not pigmented, P-mean = -2.598, 95% CI = -8.734 – 3.811, P_MCMC_ = 0.375). These results were confirmed by equivalent models constructed using phylogenetic generalized least-squares (see Table S5). No interaction between pigmentation and shape was found when human-perceived leaf masquerade presence/absence was used as the dependent variable rather than “leafiness” score, possibly because the former metric was too coarse to capture continuous variation in leafiness (MCMCglmm: P-mean = -17.816, 95% CI = -56.320 – 6.498, P_MCMC_ = 0.128; Table S4,S5). Additionally, our trichromatic human-based metrics of leafiness might represent a conservative estimate of how colour and shape interact when perceived by ecologically relevant New World primates, whose dichromatic visual systems are thought to improve the ability to find camouflaged prey [23,27,28]. Overall, in concordance with the conclusions derived from our artificial prey experiment, perception of leaf masquerade appears reliant on the co-occurrence of appropriate colouration, shape, and possibly other independent components, for exploiting both avian and mammalian recognition systems.

### Colouration and shape co-evolve synergistically to produce spectacular leaf masquerade

Our analyses indicate that colouration and shape coevolved to accelerate the evolution of masquerade as a leaf. After controlling for phylogeny, MCMCglmm analysis on individual data and a phylogenetic ANOVA on species means confirmed that the evolution of green pigmentation was significantly associated with reduced tegmen aspect ratios (MCMCglmm: P-mean = 0.061, 95% CI = 0.003 – 0.114, P_MCMC_ = 0.032; phylogenetic ANOVA: F_1_ = 16.469, p = 0.018; Figure 2D). Given this evolutionary correlation, and the composite nature of leaf masquerade, we tested the order of pigmentation, green colouration, and aspect ratio acquisition across the phylogeny (using phylogenetic pathway analysis). When both human-perceived metrics of leafiness were used as the dependent variable, our models showed strong support for the coordinated synergistic evolution of all three traits, consistent with their simultaneous acquisition during the evolution of leaf masquerade (Figure 2E; Figure S3; Table S6).

### Conclusion

Functional synergy between colour and shape of leaf-like prey causes misclassification by naïve predators, and phylogenetic reconstruction of these component traits supports their simultaneous, synergistic evolution in Neotropical katydid wings. Our field experiment demonstrates that resemblance in both colour and shape is necessary for leaf masquerade to be functional. While we isolate and demonstrate the functional benefits of masquerade in a natural context and illustrate the synergistic evolution of its component traits, other sources of selection are likely to act upon the wings of neotropical katydids, including sexual selection, as male wings are used to produce sound, and other forms of ecological selection. In nature, masquerade likely also co-evolves with behavioural adaptations such as resting orientation and microhabitat selection, as has been observed in other masquerading organisms [8,29-31]

By integrating experimental evidence with comparative analyses of extant species, we demonstrate how composite adaptations can originate through the simultaneous recruitment of individual trait components, rather than by stepwise evolution. Our findings validate and expand upon patterns of contingent wing pattern evolution observed in leaf-masquerading butterflies [32,33], providing a plausible evolutionary mechanism for explaining the origin of integrated phenotypes, leaving its genetic basis an alluring topic of future study. Although selection for background matching may have contributed towards the evolution and maintenance of leaf masquerade, disentangling relative strengths of these effects, particularly with little ecological data, is challenging. It is irrelevant to our arguments how much leaf masqueraders benefit from crypsis through background matching; their often-uncanny specific resemblance to leaves suggests an additional benefit through being misidentified. In many species, leaf masquerade appears to have been further optimised through the evolution of elaborate venation patterning, false necrotic spots, and false holes [12,31,34]. The sequence and pattern by which these more sophisticated leaf-specific traits are acquired across species, and their relative fitness advantages, warrants further investigation. The appearance of exquisitely sophisticated ‘design’ in the natural world is an enduring issue in evolutionary biology that propels both historical and contemporary debates [4,5,35]. While the dominant, gradualist paradigm envisions sequential optimisation of adaptive traits that eventually build complex functions, we suggest that alternative scenarios warrant more serious investigation; that is, those involving the co-occurrence of traits with synergistic effects, which may be rarer, but subject to stronger selection.

## Materials and methods

### Wild katydid sampling

Katydids were sampled at night from lights around research station buildings on Barro Colorado Island (BCI), Panama (9.1647° N, 79.8367° W) in March-April 2024 between 04:00 and 06:00 and 20:00 and 01:00. Individuals were identified to species level using public resources [36.37] and custom ID keys provided by Dr Laurel Symes and Dr Hannah ter Hofstede. A total of 270 individuals were sampled across 51 species, with representatives from subfamilies Conocephalinae, Phaneropterinae and Pseudophyllinae (Figure 1A,2A). We collected more than three individuals of 28 species. Studying a single diverse community of katydids with seemingly generalist habitat preferences [17] allowed for robust comparative analyses by eliminating variation in background vegetation composition, which might influence the evolution of different leaf masquerading strategies.

Captured individuals were placed in a freezer at -20 °C for ten minutes, after which morphological data were collected (body length, femur length/width, thorax height). Whole tegmina were removed with microscissors. The external surface of the tegmina was photographed using a Nikon D3300 camera (Nikon Corporation, Tokyo, Japan) with a Sigma 17-50 mm f/2.8 lens (Sigma Corporation, Kanagawa, Japan) in a photography light box (DUCLUS, model DU5032U, Guangdong, China) under LED illumination. Images were taken against a matte white background, at a constant distance (∼25 cm) and magnification, and contained a ColorChecker passport (X-Rite Inc. Grand Rapids, MI, USA) for future colouration analysis. Tegmina were preserved as dry specimens, and the remaining body tissue was either preserved in 95% ethanol or contributed to the Museo de Invertebrados Fairchild de la Universidad de Panamá (MIUP). Samples were exported to the United Kingdom under export permit collection no. PA-01-ARG-040-2024, obtained from the Ministerio de Ambiente, Panama.

### Field experiment

#### Target creation

To test whether combinations of colour and shape provide fitness benefits through masquerade in nature, artificial leaf-like stimuli were exposed to wild avian predators. Prey targets consisted of coloured paper ‘leaves’ attached to an edible mealworm ‘body’ (*Tenebrio molitor* larvae, frozen at -80°C then thawed). All treatments had a total surface area of 4.85 cm^2^ which falls within the range of a) suitable prey typically consumed by local passerine birds and b) leaf area variation observed in real bramble (*Rubus spp.*) leaves which these targets were designed to, at least somewhat, imitate [38]. Three treatment groups for both colour and shape were created, following a 3 x 3 factorial design.

To create calibrated green stimuli, digital photographs of 34 individual bramble leaves were taken using a Nikon D3300 camera (Nikon Corporation, Tokyo, Japan) with a Sigma 17-50 mm f/2.8 lens (Sigma Corporation, Kanagawa, Japan) under natural daylight illumination, each containing a ColorChecker passport (X-Rite Inc. Grand Rapids, MI, USA) for colour calibration [39]. Bramble was chosen because it was common across our field sites, meaning local birds would have encountered its leaves, but crucially, it and any other green foliage was never found on the tree bark substrates on which targets were pinned. Images were saved in raw (NEF) format and imported into ImageJ [40] for subsequent linearisation and calibration within the MICA toolbox software attachment [41]. Multispectral images were converted to blue tit (*Cyanistes caeruleus*) visual colour space, a model insectivorous passerine bird present at our field site [42]. From these cone-catch images, the mean RGB values of each leaf were obtained and averaged across all images to estimate the average colour of bramble leaves as perceived by ecologically relevant predators.

To separate the effects of masquerade from those of background matching camouflage, brown stimuli that matched the average colour of the substrate upon which targets were pinned formed a second colour treatment group. A total of 57 photographs of oak (*Quercus robur*), ash (*Fraxinus excelsior*), sycamore (*Acer pseudoplatanus*) and field maple (*Acer campestre*) bark were taken and calibrated using the same methodology as above. As a control, blue targets that did not match any aspect of the environment were created by modifying the RGB values of the green stimulus such that the overall luminance, measured in double cone catch quanta (the avian achromatic channel [41,42]), was the same.

The shape of the artificial stimuli was designed based on the mean aspect ratio of tegmina estimated from *Aegimia maculofolia* (Tettigoniidae: Phaneropterinae) (2.101; n = 7), a highly specialised leaf-masquerading katydid species [43]. Aspect ratio is an accurate proxy of leafiness where individuals with low aspect ratios have more rounded, ‘leaf-like’ wings [11,18]. The maximum length (the ‘major axis’) and width (the ‘minor axis’, perpendicular to the length) of the left and right tegmen was measured from JPEG light box images (see above) in ImageJ^37^, averaged, and used to calculate the “tegmen aspect ratio” for this species (aspect ratio = major axis length / minor axis length). The objective was to generate a shape that was recognisably leaf-like but was not under any developmental constraint, such that it could plausibly evolve in the wings of real insect prey.

A generic leaflike shape with this aspect ratio (3.72 x 1.77 cm) was made by creating a vesica piscis using the shape tool in Powerpoint (Microsoft Corporation). An elongated treatment was created by doubling the length of the original vesica piscis whilst keeping the total surface area the same. The resulting shape (7.45 x 0.88 cm) had an aspect ratio of 8.47. This aspect ratio of the elongated treatment did not resemble that of most natural leaves, allowing us to control for intrinsic Gestalt properties of being leaf shaped which could predict the detectability of prey during visual search, rather than providing a benefit through misclassification (i.e. masquerade) *per se*. In addition, this treatment evaluated whether elongate-bodied insect species (including all sampled katydids) generally benefit from masquerade to some degree. As a control, a non-leaf shaped rectangular treatment was created, with surface area kept constant (3.19 x 1.52 cm).

Pairs of adjacent targets of the same treatment group were printed double-sided onto A4 waterproof paper (Rite-in-the-Rain, JL Darling, Tacoma, WA, USA) using a calibrated Xerox VersaLink C7030 printer (Xerox, CT, USA). These were then folded along the midline (where the boundaries of the adjacent pairs overlapped slightly), with a steel sewing pin (Korbond Industries Ltd. Lincolnshire, UK) driven through the centre, and glued together (UHU, Bühl, Germany).

#### Protocol

The field experiment took place from July to September 2024 and was conducted along the Lade Braes public footpath, St Andrews, Fife, UK (56.3371° N, 2.8036° W), a mixed-deciduous forested habitat, covering an area of approximately ∼1.7 km^2^, and inhabited by a variety of insectivorous passerine birds. Replicate blocks of targets were pinned at three self-contained, unfragmented sites along this footpath, which ran parallel to the Kinness Burn (Cockshaugh Park site) and the Cairnsmill Burn (eastern and western sites). The experiment was approved by the School of Biology Ethics Committee, University of St Andrews (reference number BL17958).

The experimental procedure followed that of several preceding studies [34,44,45]. Targets were haphazardly selected from a plastic bag in which all targets for a block had been thoroughly mixed. These were then pinned to the bark of oak, ash, sycamore and field maple exclusively (to ensure brown targets matched the colour of their background) with a thawed mealworm threaded onto the pin (Figure 1B). Targets were pinned to the bark of tree trunks free of lichen, moss, or vegetation, at a height of approximately 1.8 m, and trees with a circumference smaller than 0.5 m were ignored. A total of 1,296 individual targets were put out across 16 experimental blocks of 81 models (nine replicates per treatment per block). Each block was conducted at a different location within each site to minimise the probability of the targets being encountered by the same individual predators. Fresh targets were made for each block.

Checks were made at 24, 48 and 72 h intervals where the ‘survival’ of an individual target at each check was determined by the presence or absence of the mealworm bait, with the target still present, intact and attached to the tree. Targets were classified as ‘censored’ if they survived until the final 72-hour check, showed signs of non-avian predation (e.g. hollow exoskeleton from spiders or slime trails from slugs), were lost, or were relocated but no longer attached to the tree. Predated targets were removed at each check, and all remaining targets were collected at the final 72-hour check.

#### Survival analysis

Survival analysis was performed by constructing mixed effects Cox regressions with the coxme() function from the *coxme* package in R [46,47]. Target colour, shape, and their interaction were treated as fixed effects, with experimental block and site included as random effects. Visual inspection of the partial residuals against the ranked survival time confirmed the proportional hazard assumption of the models. Analyses of deviance, comparing models with and without the factor of interest, were then performed using the anova() function and tested against a χ^2^ distribution. Subsequent post-hoc pairwise contrasts were made by creating custom contrasts using the glht() function from the *multcomp* package [48]. P-values were unadjusted if the number of pairwise tests was not greater than the degrees of freedom [49].

### Comparative morphology

#### Katydid wing traits

From the field-collected katydid tegmen samples, colour and shape data were obtained to investigate how these traits coevolve to promote the elaboration of leaf-masquerading phenotypes across species. After randomly sampling a single light box image per species, a researcher scored whether the majority of the tegmen surface was a) pigmented or not and b) green or brown in colour (Figure 2B). In ‘unpigmented’ species, the cells created by the reticulations of the tegmen venation were clearly visible, indicating a lack of pigment in these regions, as they appeared either translucent or completely transparent. Both colouration classifications were confirmed independently by a naïve researcher who scored the same images and had not been informed about the purpose of the study. Based on these classifications, species were then grouped based on whether they were green *and* pigmented or not.

To ensure accurate measures of shape, a different set of images were taken using flattened tegmen samples. Dried tegmina were rehydrated in an insect relaxing chamber: a hermetic plastic box filled with paper towels soaked in water, with a small quantity of ethanol to prevent contamination, for 24 hours. This procedure softened the cuticle, easing manipulation of the sample by reducing the risk of shattering. Individual tegmina were then mounted between two clean microscope slides and sealed with scotch tape. Whole specimens were photographed, accompanied by a suitable scale, with the tripod-mounted 12MP camera of an iPhone 13 Mini (Apple Inc., Cupertino, CA, USA) under ventral LED illumination. JPEG images were imported into ImageJ [40] where the tegmen aspect ratio for each individual was calculated (Figure 2C). The length of the minor axis was measured excluding the stridulatory area which naturally forms a perpendicular fold that rests on the dorsal surface of the animal. Mean aspect ratio for each species was calculated, and both individual-level and mean values were log_10_ transformed prior to analysis.

To explore how colour and shape influence human perceptions of leafiness, an online survey was designed and distributed using Qualtrics (Qualtrics Provo UT, USA) where a randomly selected light box image of each katydid species was shown to 53 participants naïve to the purpose of the study (53% female, aged 18+, all with trichromatic colour vision). All participants gave their informed consent in line with the Declaration of Helsinki. For each participant, the 51 images were presented in a randomised order. Participants were given one minute per image to a) state whether each species was ‘leaf-like’ or not and b) provide a ‘leafiness score’ of each species on a 0-10 scale, where 0 is not leaf-like at all and 10 is extremely leaf-like. Once a response was submitted for each species, participants were not able to change their mind. The survey was approved by the School of Biology Ethics Committee, University of St Andrews (reference number BL18128).

#### Phylogenetic comparative analysis

Using a recent phylogenetic tree for Tettigoniidae from Kernan *et al.* [19], pruned to include only the 51 sampled species, the phylogenetic signal of each trait was estimated using Pagel’s λ [50]. To model the evolution of binary colouration traits, the fitDiscrete() function from the *geiger* package was implemented [51]. The equal rates (ER) model (where the probability of transitioning into each character state is equal) was compared with the all-rates-different (ARD) model (where the probability of transitioning into each character state is unequal) based on the difference in Akaike’s information criterion (ΔAIC). Models with a ΔAIC < 2 are considered equivalent and therefore, the model with the fewest parameters (ER) is selected. The best fitting model was used to run the make.simmap() function in *phytools* [52] which applies a maximum likelihood approach to simulate 10,000 character maps of the phylogeny and reconstructs the evolution of colouration across the tree (Figure S2). This provides an estimate of the colouration state at each node, including the ancestral character state.

Aspect ratio was modelled by constructing Brownian motion (BM; random-walk evolutionary processes with no selection component), Ornstein-Uhlenbeck (OU; stochastic variation with a deterministic selection component), and early-burst (EB; rapid diversification early in cladogenesis followed by Brownian motion) models of trait evolution using the fitContinuous() function in *geiger* [51]. As with colouration, models were compared based on ΔAIC and visualised using the contMap() function [49] (Figure S2).

To test if colouration, mean aspect ratio, and their interaction influence human perceptions of leafiness, Bayesian phylogenetic generalized linear mixed models were constructed using the *MCMCglmm* package in R, which incorporates the inverse correlation matrix of the phylogeny as a random effect [53]. Default priors were used as fixed effects. For normally distributed continuous (family = “gaussian”) response variables, uninformative, parameter expanded priors were used as random effects (G: V = 1, nu = 1, alpha.mu = 0, alpha.V = 1,000; R: V = 1, nu = 0.002), whereas with binary (family = “categorical”) and ordinal response variables (i.e. Leafiness score; family = “ordinal”), weakly informative inverse-Wishart priors were used with a fixed residual variance (binary variables, G: V = 1, nu = 1, alpha.mu = 0, alpha.V = 1000; R: V = 1, nu = 0.002, fix = 1; ordinal variables, G: V = 1, nu = 0.002; R: V = 1, fix = 1). Species and participant were included as additional random effects and each model was run for 5,100,00 iterations, with a burnin of 100,000. Additional models were also run to test for evolutionary associations between individual-level aspect ratio and colouration. In these models, only species was included as a random effect. In all cases, the significance of each predictor is reported as the probability of the parameter value being different from zero (*P_MCMC_*). We also report the posterior mean (P-mean) and 95% credible interval for each predictor.

By calculating the proportion of participants categorising each species as leaf-like and the mean leafiness score per species, equivalent models to those made with MCMCglmm were recreated using phylogenetic generalized least-squares regression (PGLS) in the *caper* package [53]. Visual inspection of the residuals from the fitted models, followed by Shapiro-Wilk tests, confirmed they were normally distributed with no heteroscedasticity. Pagel’s λ for each model was estimated based on maximum likelihood. To test for an association between mean aspect ratio and colouration, phylogenetic ANOVAs were performed, under 1000 simulations, using the phylANOVA() function in the *phytools* package [52].

Finally, we modelled support for hypothetical causal relationships between traits using phylogenetic pathway analysis (PPA) with the *phylopath* package [55,56]. This method integrates categorical (colouration) and continuous (mean tegmen aspect ratio) trait data to assess the interrelation between traits that result in “better” leaf masquerade, based on the human-informed metrics of leafiness described above. By building structural equation models that suggest multiple scenarios by which colouration and shape are acquired during leaf masquerade evolution (Figure S3), the best fitting causal structure is indicated by the model with the lowest C-statistics information criterion (CIC).

## Acknowledgements

We are grateful to Gregg Cohen and Rachel Page at the Smithsonian Tropical Research Institute, all staff at the Barro Colorado Island research station, Lil Marie Camacho, and Panama’s Ministerio de Ambiente for logistical fieldwork support and Laurel Symes for help with katydid identification. Many thanks also to Innes Cuthill and Jolyon Troscianko for providing advice on colour calibration, Rivers Tarr and Sophie Rolfe for helping with target creation, and Mirabella Funk for assistance in the field. Lastly, we thank Tony Robillard for providing permission to use his phylogenetic tree, Leeban Yusuf for help with colouration categorisation, and all participants involved in the online survey; they were not financially remunerated.

## Data availability

All raw data and analysis R code have been deposited at Zenodo alongside thecalibrated targets used in the field experiment, all lightbox images used in the online survey and mounted tegmen images (https://tinyurl.com/35xc343f) and will be publicly available as of the date of publication.

## Supplementary figures and captions

**Figure S1.**
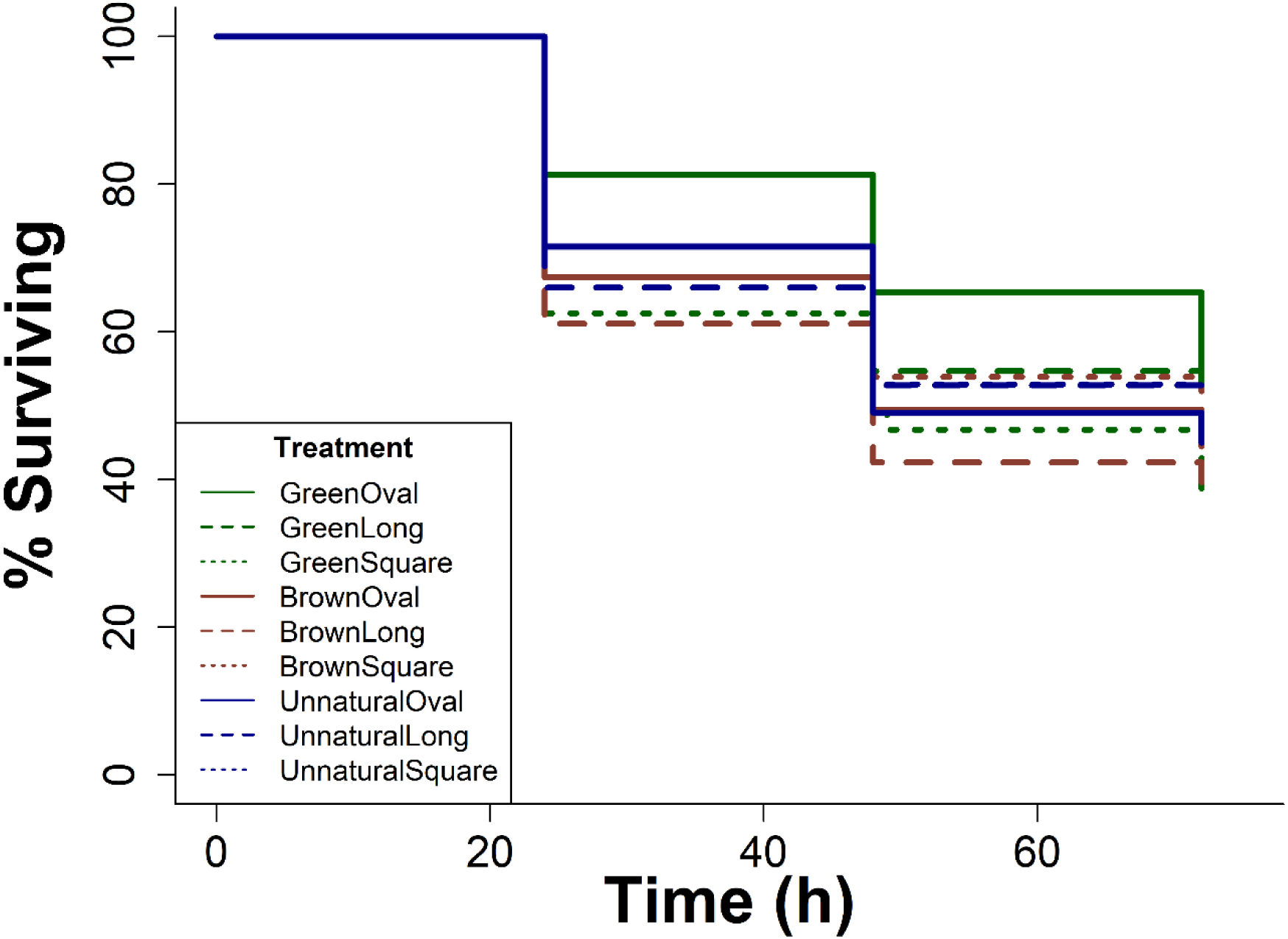
Survival plot for each colour x shape treatment combination over time when exposed to wild avian predation (*N* = 1,296).

**Figure S2.**
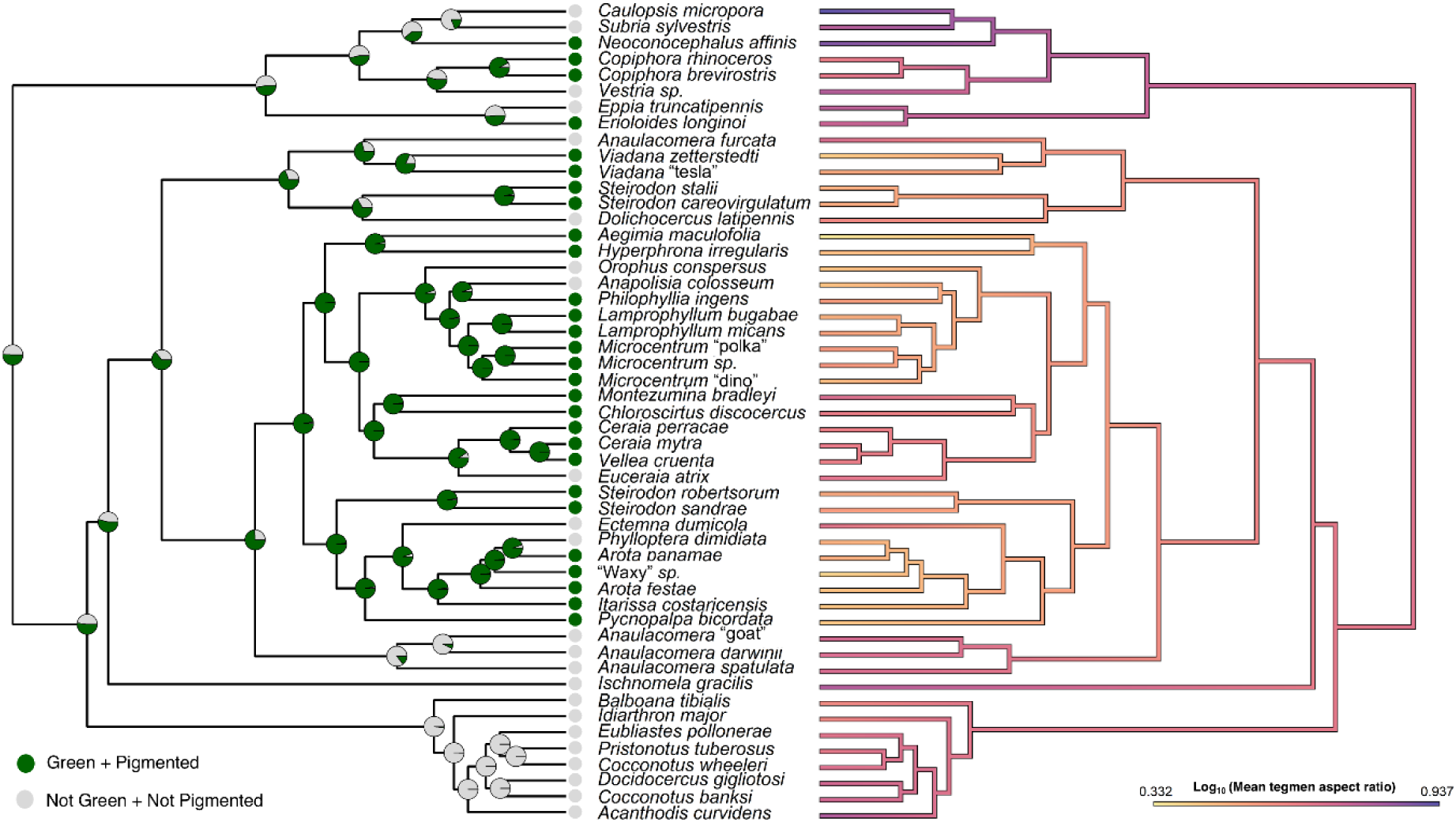
Estimated transitions from absence (grey nodes and tips) to presence (green nodes and tips) of green tegmen pigmentation, and *vice versa* (left), and patterns of mean tegmen aspect ratio (au) evolution (right) across a pruned molecular phylogeny of 51 sampled katydid species [19].

**Figure S3.**
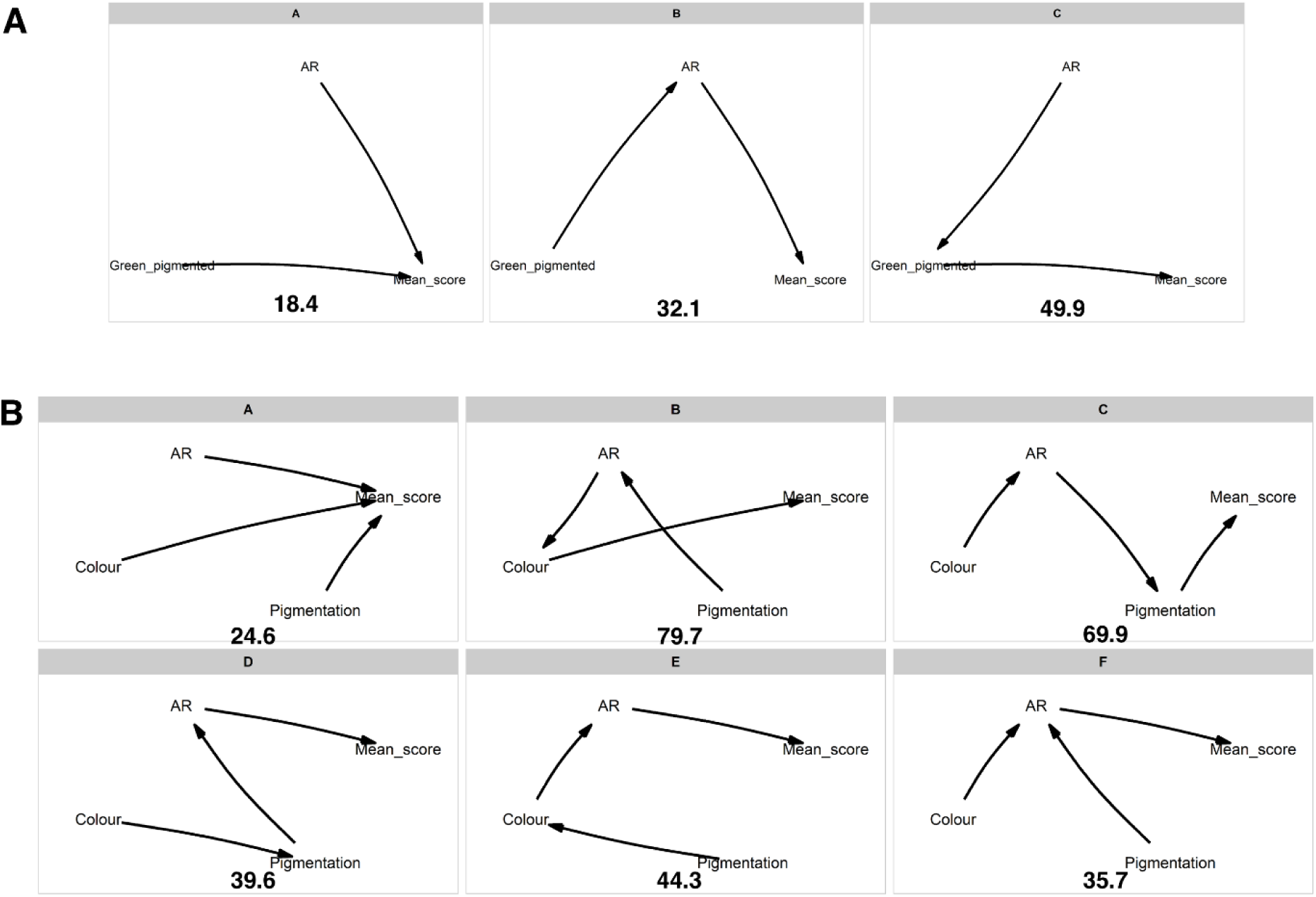
All models included in each set of the phylogenetic pathway analysis with mean leafiness score as the response variable. The C-statistics information criterion (CIC) is indicated below each model. (A) Model set included the presence or absence of green pigmentation (Green_pigmented) and mean tegmen aspect ratio (AR) as predictor variables. (B) Model set included mean tegmen aspect ratio and the presence or absence of pigmentation, and green colouration (‘Colour’) as individual predictor variables.

## Supplementary tables and captions

**Table S1.** Results from post-hoc custom pairwise comparisons testing the effect of colour on the survival of oval shaped targets in the wild avian predation experiment, the only shape treatment where colour had a significant effect. The survival rate of the ‘green’ treatment was compared with that of the ‘brown’ and ‘blue’ colour treatments. Significant differences are denoted by asterisks. *p < 0.05, **p < 0.01, ***p < 0.001.

| Pairwise treatment contrast | Estimated coefficient | Standard error | z value | p value |
| --- | --- | --- | --- | --- |
| ‘Green oval’ – ‘Brown oval’ | -0.498 | 0.203 | -2.456 | 0.014* |
| ‘Green oval’ – ‘Blue oval’ | -0.438 | 0.209 | -2.090 | 0.037* |

**Table S2.**
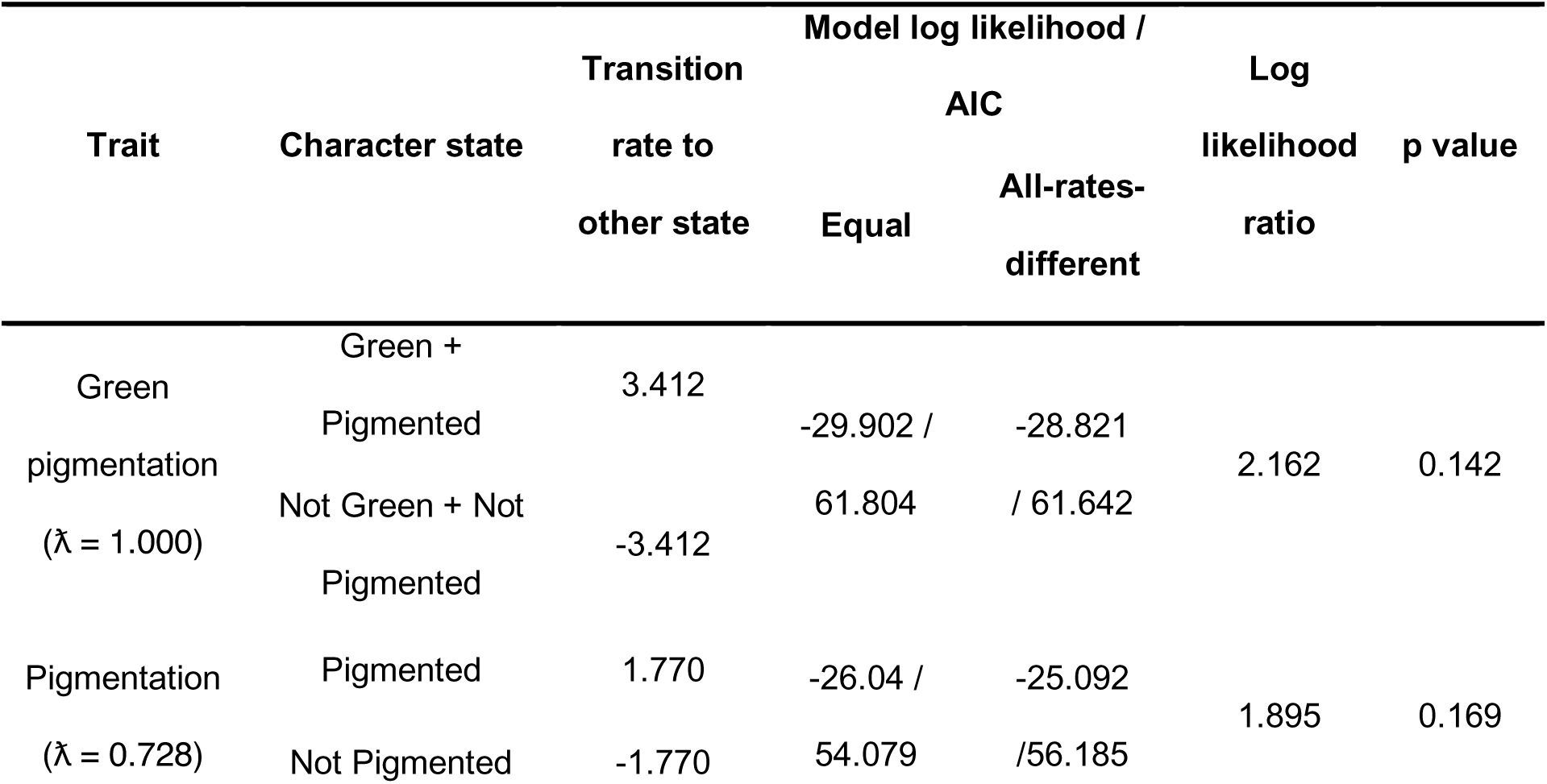
Transition rates between colouration character states across the katydid phylogeny. Included are the log likelihood and AIC values of the equal rates (null) model and the all-rates-different model. Output from log likelihood ratio tests, comparing the two models are also shown where a significant difference (p < 0.05) confirms that the all-rates-different model is accepted.

**Table S3.** Evolutionary modelling of continuous traits across the katydid phylogeny. Included are log likelihood and AIC values for Brownian motion (BM), Ornstein-Uhlenbeck (OU), and early burst (EB) models. Output from likelihood ratio tests, comparing each model with the one with the lowest AIC score are also shown.

| Trait | Model log likelihood / AIC |  |  | Log likelihood ratio / p value |  |
| --- | --- | --- | --- | --- | --- |
|  | BM | OU | EB | BM vs. OU | BM vs. EB |
| Mean tegmen aspect ratio<br>($\lambda = 1.061$) | 49.650 / -95.050 | 49.650 / -92.789 | 49.925 / -93.340 | 0.000 / 1.000 | 0.551 / 0.458 |
| Leaf masquerade probability<br>($\lambda = 0.905$) | 5.577 / -6.904 | 6.363 / -6.216 | 5.577 / -4.644 | 1.572 / 0.210 | 0.000 / 1.000 |
| Mean leafiness score<br>($\lambda = 1.002$) | -92.155 / 188.561 | -91.531 / 189.572 | -92.155 / 190.822 | 1.250 / 0.264 | 0.000 / 1.000 |

**Table S4.**
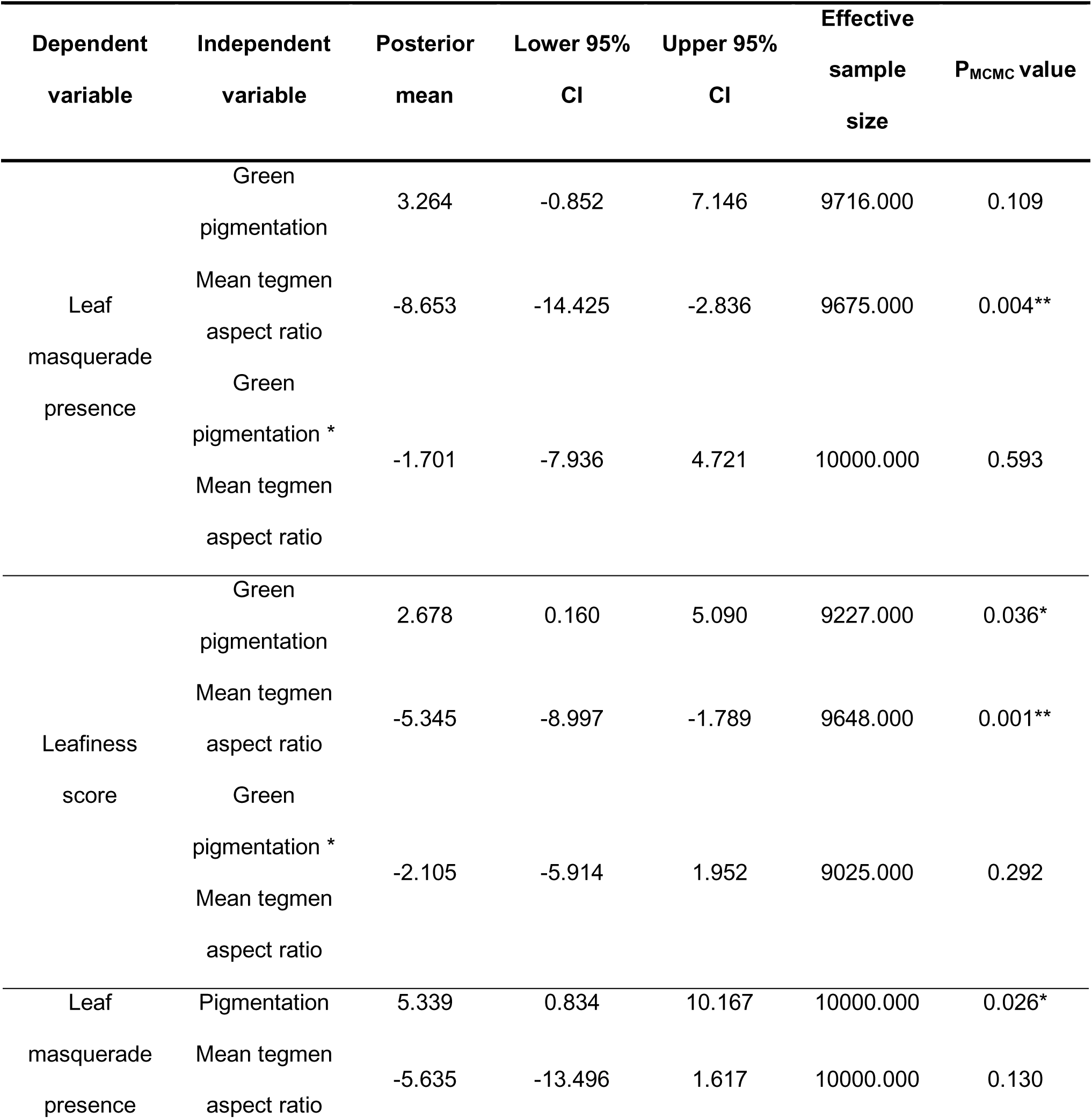
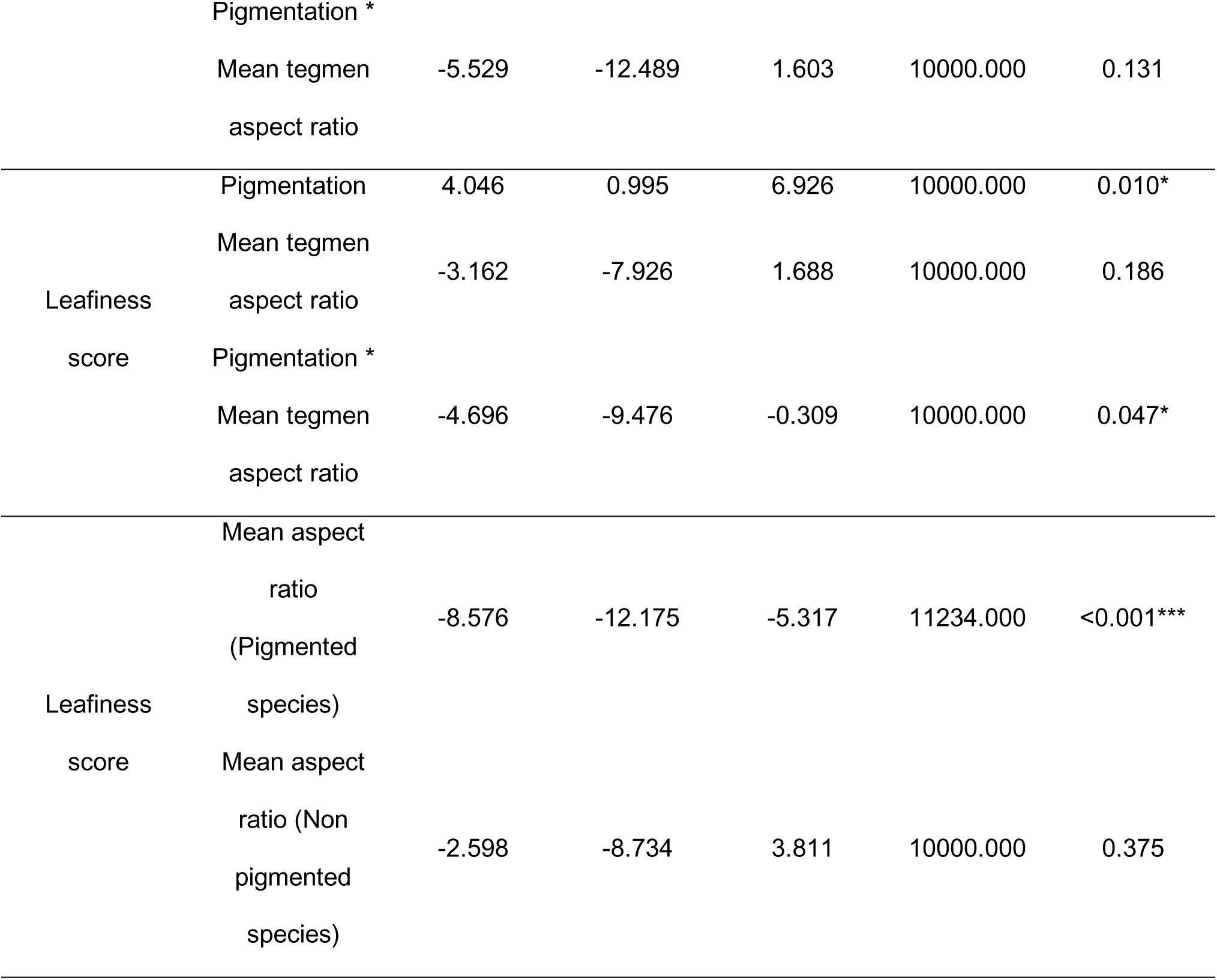
Results from MCMCglmms which test the effect of colouration (green pigmented, pigmented), shape (mean tegmen aspect ratio) and their interaction on individual human leafiness perception (Leaf masquerade presence and Leafiness score). Shown are the posterior means, 95% credible intervals, and the effective sample size for each independent variable in the model. Significant effects are denoted by asterisks. *P_MCMC_ < 0.05, ** P_MCMC_ < 0.01, *** P_MCMC_ < 0.001.

**Table S5.**
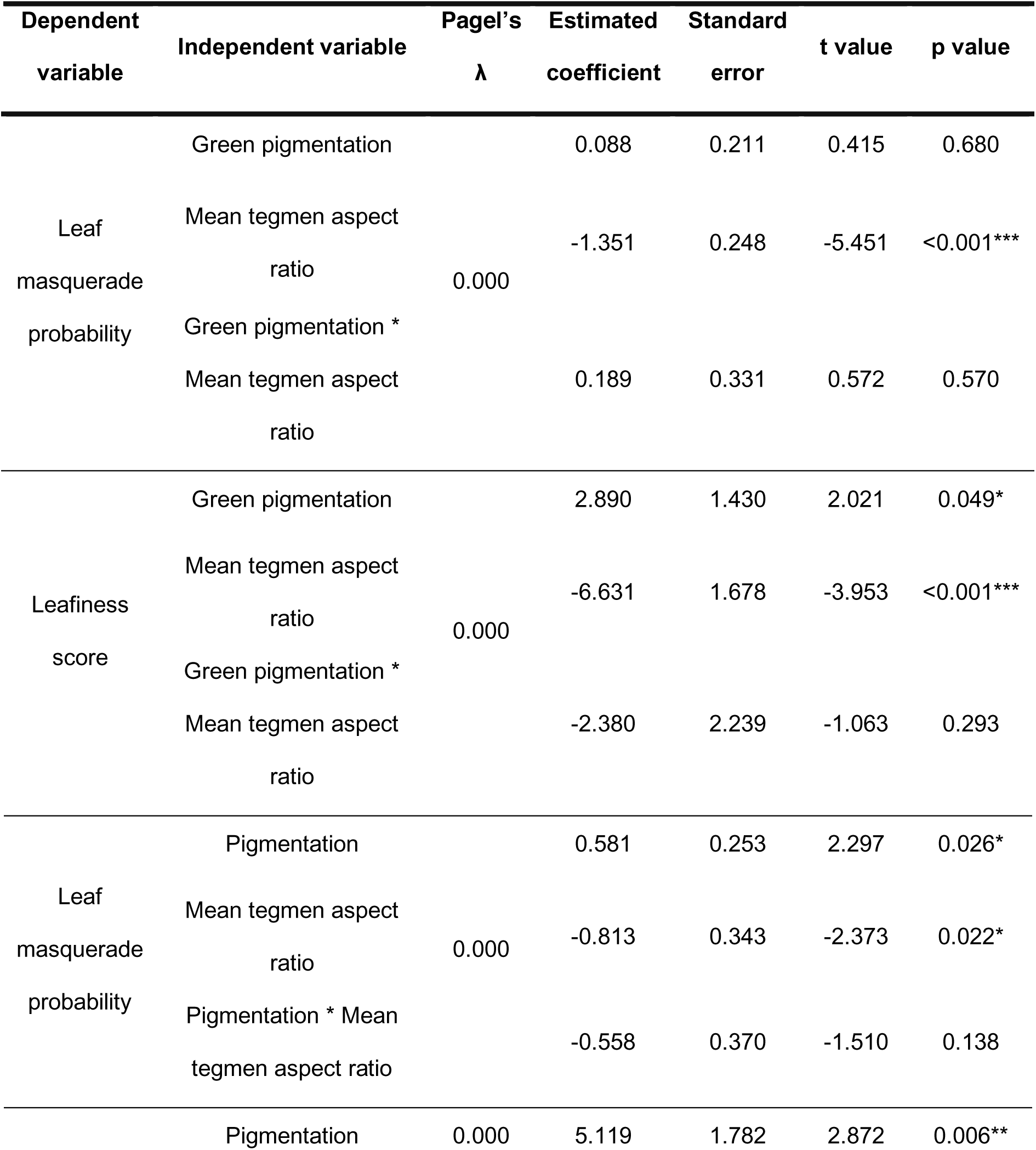
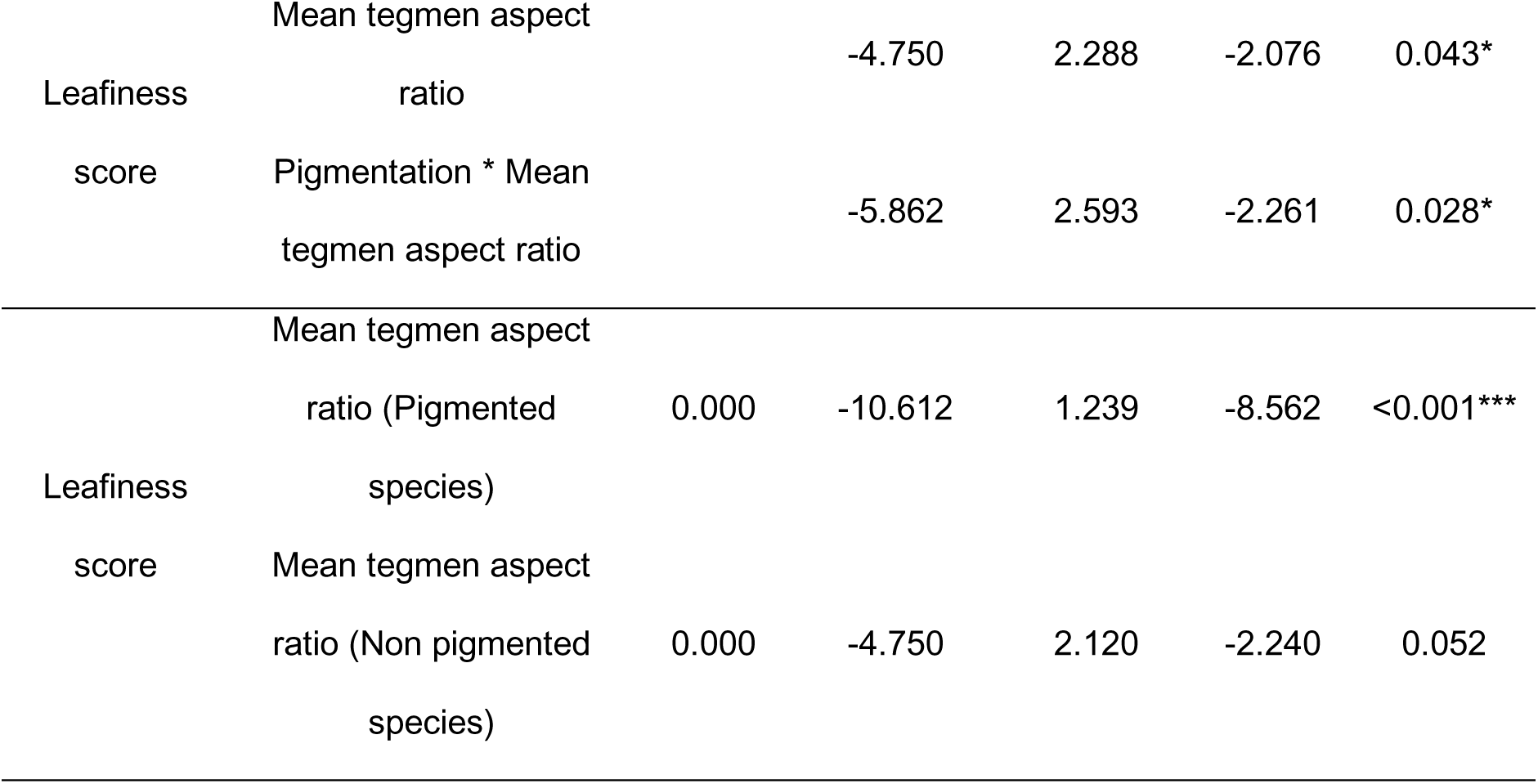
Results from phylogenetic generalized least-squares (PGLS) models which test the effect of colouration (green pigmented, pigmented), shape (mean tegmen aspect ratio) and their interaction on the species means of two metrics of human leafiness perception (Leaf mimicry probability and Mean leafiness score). Phylogenetic signal of each model was calculated using the maximum likelihood estimate of Pagel’s λ. Significant effects are denoted by asterisks. *p < 0.05, **p < 0.01, ***p < 0.001.

**Table S6.** Pathway coefficients from the best fitting model from four separate phylogenetic pathway analysis model sets, alongside the standard error and 95% confidence intervals.

| Model set | Pathway | Path coefficient | Standard error | Lower 95% confidence interval | Upper 95% confidence interval |
| --- | --- | --- | --- | --- | --- |
| Green pigmentation (GP), mean tegmen aspect ratio (AR), leaf masquerade probability (LP) | GP -> LP | 0.746 | 0.162 | 0.433 | 1.106 |
|  | AR -> LP | -0.622 | 0.081 | -0.775 | -0.458 |
| Green pigmentation (GP), mean tegmen aspect ratio (AR), leafiness score (LS) | GP -> LS | 0.793 | 0.164 | 0.482 | 1.127 |
|  | AR -> LS | -0.601 | 0.082 | -0.752 | -0.449 |
| Colour (C), Pigmentation (P), mean tegmen aspect ratio (AR), leaf masquerade probability (LP) | C -> LP | 0.543 | 0.163 | 0.214 | 0.860 |
|  | P -> LP | 0.824 | 0.175 | 0.510 | 1.162 |
|  | AR -> LP | -0.652 | 0.074 | -0.794 | -0.523 |
| Colour (C), pigmentation (P), mean tegmen aspect ratio (AR), leafiness score (LS) | C -> LS | 0.569 | 0.187 | 0.248 | 0.919 |
|  | P -> LS | 0.823 | 0.197 | 0.467 | 1.169 |
|  | AR -> LS | -0.613 | 0.084 | -0.761 | -0.464 |

